# Low-dimensional neural dynamics underlying rhythmic vocal behavior in songbirds

**DOI:** 10.64898/2026.03.28.715006

**Authors:** Fiamma L. Leites, Cecilia T. Herbert, Santiago Boari, Felipe I. Cignoli, Gabriel B. Mindlin, Ana Amador

## Abstract

Birdsong is a complex learned behavior that requires millisecond-scale precision in the coordinated activation of respiratory and vocal muscles to generate sound. Canary song consists of sequences of syllables organized into phrases, in which each syllable type is repeated at a characteristic rate, giving rise to a well-defined rhythmic vocal behavior. Here, we analyze neural population activity in the telencephalic song system nucleus HVC of singing adult male canaries (*Serinus canaria*), in relation to both vocal output and the underlying respiratory motor gestures. To uncover structure in these high-dimensional neural recordings, we used an unsupervised autoencoder. We found that a three-dimensional latent space was sufficient to reconstruct the data with minimal information loss, revealing a low-dimensional representation of HVC population activity. The oscillation frequencies of the latent modes closely matched both the syllabic repetition rate and the corresponding respiratory motor patterns.

These results show that multiunit activity in HVC captures key rhythmic features of song at the population level, providing a dynamical representation of behaviorally relevant motor structure. More broadly, our findings highlight how data-driven dimensionality reduction can reveal structured, low-dimensional neural dynamics underlying complex learned motor behaviors.

## Introduction

Understanding how complex learned behaviors are represented in large-scale neural population activity remains a central challenge in systems neuroscience. Temporally structured vocal behaviors, such as birdsong or speech, provide a powerful framework to address this question, as their production requires precise coordination of distributed neural populations across brain areas. Yet the high dimensionality and variability of neural recordings make it difficult to extract interpretable dynamical structure linked to behaviorally relevant features [Cunningham & Yu, 2014; Churchland et al., 2012]. Recent advances in machine learning and latent-variable modeling offer promising approaches to this challenge, enabling the identification of low-dimensional and potentially interpretable structure in large-scale neural activity [Pandarinath et al., 2018; Whiteway & Butts, 2019; Raut et al., 2025].

In this context, songbirds have emerged as a powerful model for investigating how neural population dynamics support learned vocal behavior. Songbirds possess a well-defined set of interconnected forebrain nuclei, collectively known as the song system, that are specialized for learning, production, and sensorimotor control of song. Within this circuit, the telencephalic nucleus HVC plays a critical role in song production and sensorimotor integration [Hahnloser et al., 2002; Amador et al., 2013; Hamaguchi et al., 2016]. In adult male canaries (*Serinus canaria*), HVC exhibits complex neural dynamics during singing [Alonso et al., 2015, 2016; Herbert et al., 2020; Cohen et al., 2020], while producing a rich and flexible repertoire of syllables with specific properties [Alliende et al., 2010; Del Negro, 2001; Lassa Ortiz et al., 2019]. Canary song exhibits marked flexibility across breeding seasons and natural social contexts [Leitner et al., 2001; Alcami et al., 2025]. Despite this behavioral flexibility, canary song exhibits a robust internal organization into phrases composed of repeated renditions of a given syllable type, each produced at a characteristic repetition rate (Figure 1, panels 1-2). Different phrases occur with variable probabilities across individuals, giving rise to distinct song repertoires that may change across seasons [Del Negro, 2001; Lehongre et al. 2008; Markowitz et al., 2013]. The syllabic production rate within each phrase establishes its characteristic rhythmic structure, providing a well-defined temporal framework for investigating how neural population dynamics codes behavior. Previous work has shown that neural oscillations in HVC are locked to the temporal structure of birdsong in anesthetized canaries during auditory playback of the bird’s own song [Boari et al., 2022]. These findings showed that HVC encodes rhythmic features of song under passive listening conditions. However, it remains unclear how such population-level dynamics relate to the active generation of song.

**Figure 1.**
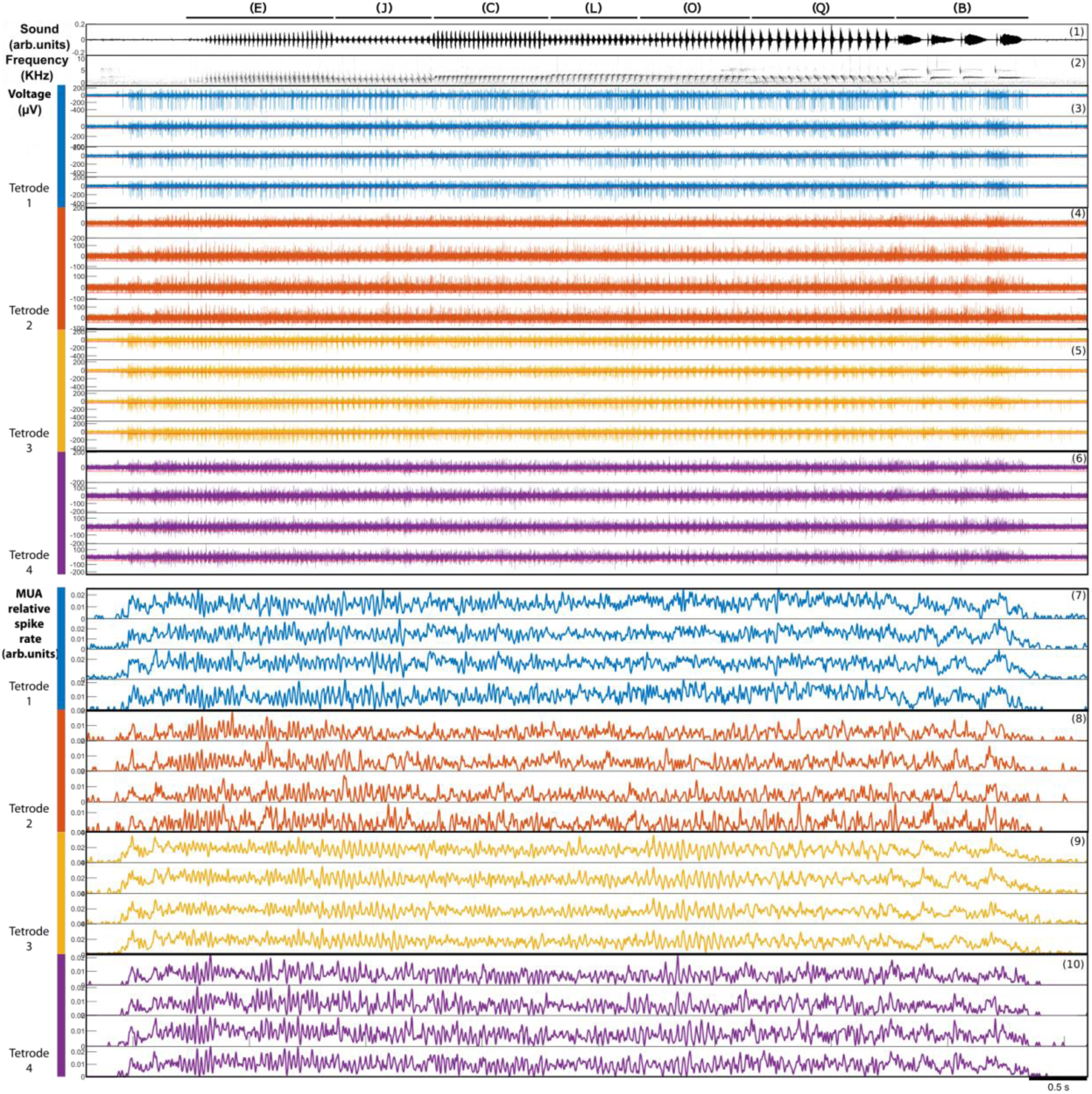
Neuronal activity during song production in canaries. Panel 1-2: oscillogram and spectrogram of a canary song. A typical canary song consists of repeating minimal units (syllables) forming phrases sung in a certain probabilistic order. Different phrases are indicated with bars and a bracketed letter. **Panels 3 to 6:** extracellular neuronal activity recorded during the song. The red horizontal line indicates the threshold used to extract the neuronal activity pattern from a population of neurons by detecting the spike times. Each color indicates a different tetrode. The detected times are used to construct the traces in panels 7 to 10 (for more details, see Methods). **Panels 7 to 10:** smoothed traces of the spiking data after using a kernel density estimation (normal kernel of bandwidth 5ms, timestep 0.033ms). These traces capture the patterns of increased activity and silences observed in panels 3 to 6. Example bird: caCTH2447-RoNa.

Although previous work has provided important insights into single-neuron activity during song production, much less is known about how HVC population dynamics encode broader temporal features during active singing. To address this, we analyze multi-unit activity (MUA) recorded in HVC of singing adult male canaries. By capturing the aggregate activity of local neuronal ensembles, MUA enables characterization of motor-related population patterns. To uncover structure in these high-dimensional population signals, we implemented an unsupervised dimensionality reduction approach based on autoencoder neural networks. By training the network to reconstruct MUA activity while compressing it into a low-dimensional latent space, we aimed to identify coordinated dynamical modes that reflect behaviorally relevant temporal structure. This data-driven strategy allows extraction of interpretable population-level dynamics directly from large-scale neural recordings, without imposing predefined features or assumptions about the underlying circuitry. This strategy allows us to address whether population-level neural activity carries an explicit signature of song rhythm during active production.

## Results

We studied multi-unit neuronal activity in HVC during song production in adult canaries. Electrophysiological recordings were obtained at multiple stereotaxic coordinates over several days in four birds. In two of these birds, air sac pressure recordings were performed on separate days following the neural recordings. Spikes were extracted from the 16 channels (four tetrodes) recorded at each stereotaxic coordinate using a threshold (-3σ; red line in Figure 1, panels 3–6) to capture events above background noise (see Methods). A smoothing procedure was applied to obtain multi-unit activity patterns (Figure 1, panels 7-10; see Methods), generating continuous traces that reflect population activity at each HVC coordinate during singing. Because signals recorded from electrodes within the same tetrode were highly similar (as the four electrodes are tightly bundled), we averaged the signals across electrodes within each tetrode. This procedure yielded four averaged traces per recording site, hereafter referred to as MUA traces (Figure 2).

**Figure 2.**
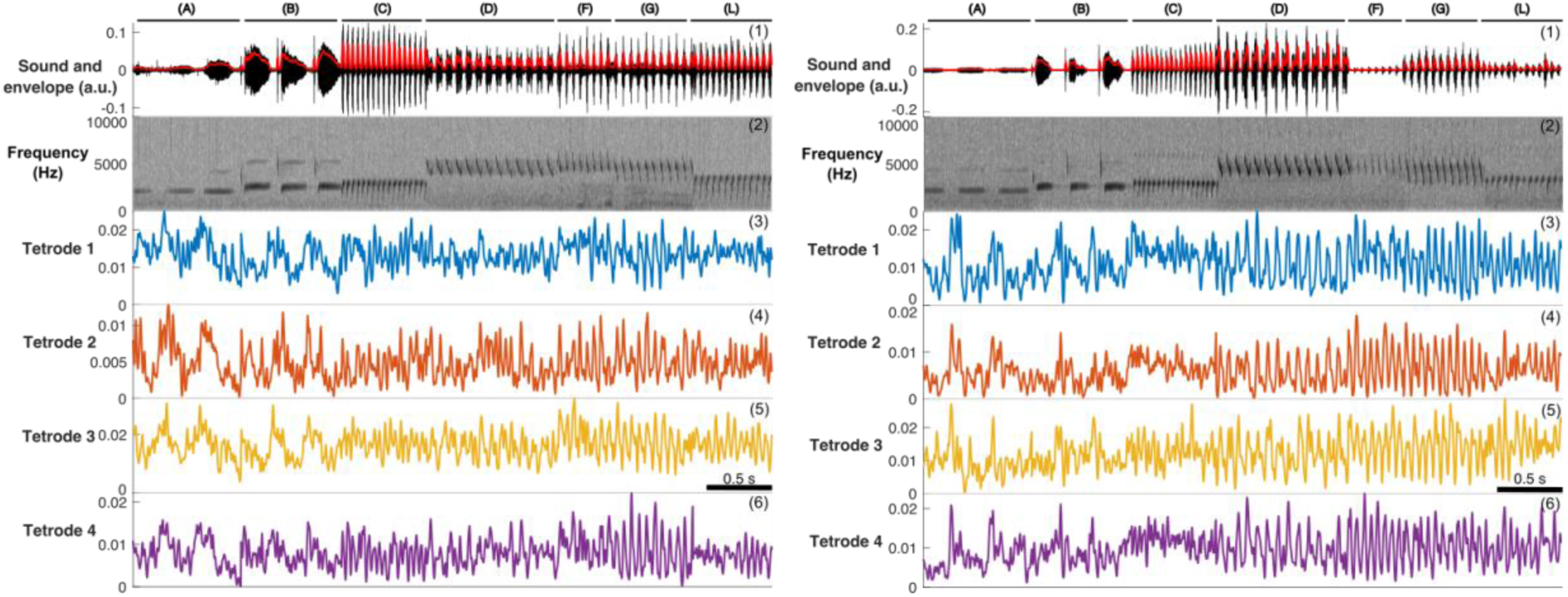
Canary song recordings and neural MUA traces (inputs for the autoencoder network). Examples of two recordings made in different stereotaxic coordinates are shown. **Panel 1:** oscillogram of a concatenated canary song, joining the selected phrases extracted from various songs performed throughout a singing session, in a fixed order for all stereotaxic coordinates. The sound envelope calculated, normalised by its mean and standard deviation, is shown in red. **Panel 2:** spectrogram of the concatenated song. **Panels 3 to 6:** MUA traces calculated for each tetrode (result of averaging the smoothed spike time traces of each tetrode, figure 1, panels 7-10), corresponding to each phrase and concatenated in the same way. Example bird: caCTH2447-RoNa.

In canaries, a phrase consists of repeated renditions of the same syllable type produced at a characteristic rate (phrases are indicated with black bars and letters in the figures). For each bird, we identified phrases in its repertoire that were sung at all recorded stereotaxic coordinates. Because our analyses focused on the rhythmic structure within each phrase, rather than on the sequential ordering of phrases in natural song, phrase segments were treated independently. Using the recorded audio as a temporal reference, we segmented each selected phrase and extracted the corresponding time-aligned MUA traces from the neural recordings. For each phrase identity, the segment duration was defined according to the minimum duration observed across recording days, ensuring consistent windowing across sites. The resulting audio and MUA segments were then concatenated to generate continuous input signals for the autoencoder. The order of concatenation was arbitrary, as the analyses were performed at the level of individual phrases and did not depend on their natural sequential arrangement. Although phrases were produced on different days, their acoustic structure was highly stereotyped, with only minor variations.

Following the processing steps described above, the resulting MUA traces revealed oscillatory activity compatible with the syllabic repetition rate in several phrases (Figure 2). However, the signals were variable across recording sites and exhibited complex structure and substantial noise, limiting the interpretability of individual traces. To extract robust population-level dynamics from these high-dimensional and heterogeneous recordings, we therefore adopted a data-driven dimensionality reduction approach based on autoencoder neural networks.

Autoencoders are neural network models trained to reconstruct their input while compressing the information into a low-dimensional latent representation. To analyze population neuronal activity recorded in HVC during singing, we trained an autoencoder to learn a compact representation of the concatenated MUA traces. The architecture of the network is shown in Figure 3A. The size of the input layer (N_input_) corresponded to the number of recording sites. Each unit consist of the temporal evolution of MUA while generating the song (Figure 2, panels 3-6). To determine the appropriate dimensionality of the latent space, we trained models with 1 to 6 latent units and evaluated reconstruction performance using the mean squared error (MSE) between input and output. As shown in Figure 3B, three latent dimensions were sufficient to achieve stable reconstruction, with no qualitative improvement in MSE beyond this point. Thus, the population activity could be accurately represented within a three-dimensional latent space.

**Figure 3.**
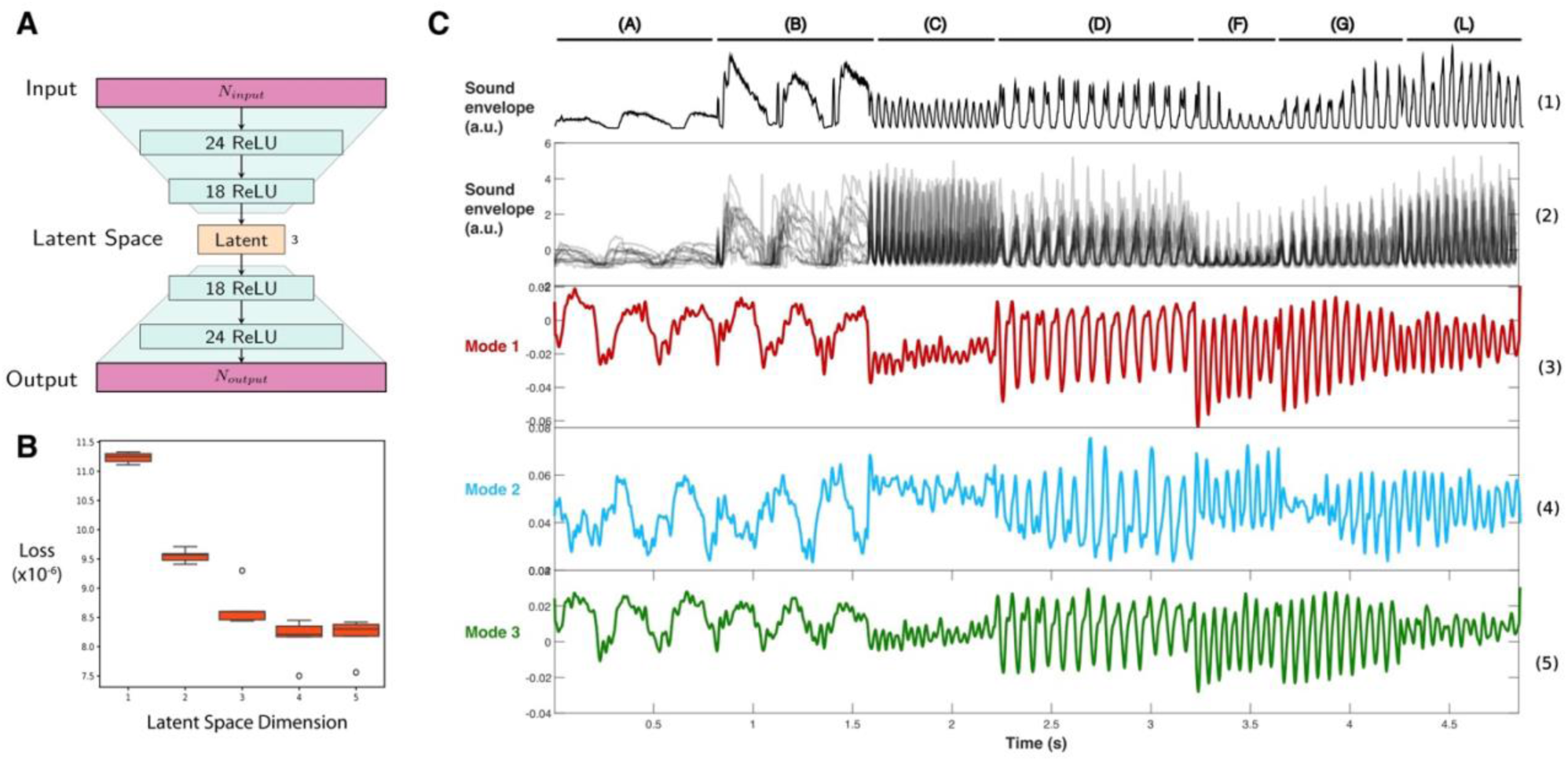
Processing of neuronal activity during canary singing using autoencoder networks. **A)** Architecture of the autoencoder network used. Each neuron was modelled as a ReLU unit, except for the neurons in the middle layer which were modelled as linear units. **B)** The dimensionality of the latent space was chosen as the minimum at which an appreciable qualitative change in the MSE is observed. From dimension 3 onwards, the decay of the mean squared error remains constant, being qualitatively lower compared to using 𝑁𝑚𝑖𝑑𝑑𝑙𝑒=2. **C)** Modes of the latent layer. Panel 1 shows an example of the sound envelope and panel 2 shows all normalized sound envelopes from all recorded stereotaxic coordinates used as input to the network for an example bird (caCTH2447-RoNa). The darker areas correspond to moments where the traces overlap, showing that while there are slight rhythmic and amplitude variations, the song is highly stereotyped. Panels 3 to 5 show the three modes of the autoencoder network’s latent layer. Example bird: caCTH2447-RoNa.

An example of the three latent modes is shown in Figure 3C (panels 3–5). Compared to the original MUA traces (Figure 2, panels 3–6), the modes exhibit more regular oscillatory structure with reduced variability. We quantified the oscillation frequencies of the three modes for each bird and compared them with the corresponding frequencies derived from the sound envelope and air sac pressure to assess whether the latent dynamics reflect behavioral rhythms. To estimate syllabic rates, we identified a consistent temporal event within each syllable across all available phrases (see Methods) and computed the inverse of the interval between successive events. An example is shown in Figure 4A, with the corresponding data points highlighted in Figures 4D and 4E. Mean values and standard errors were calculated for each phrase. Oscillation frequencies of the latent modes were estimated using the same procedure (Figure 4A, middle panel).

**Figure 4.**
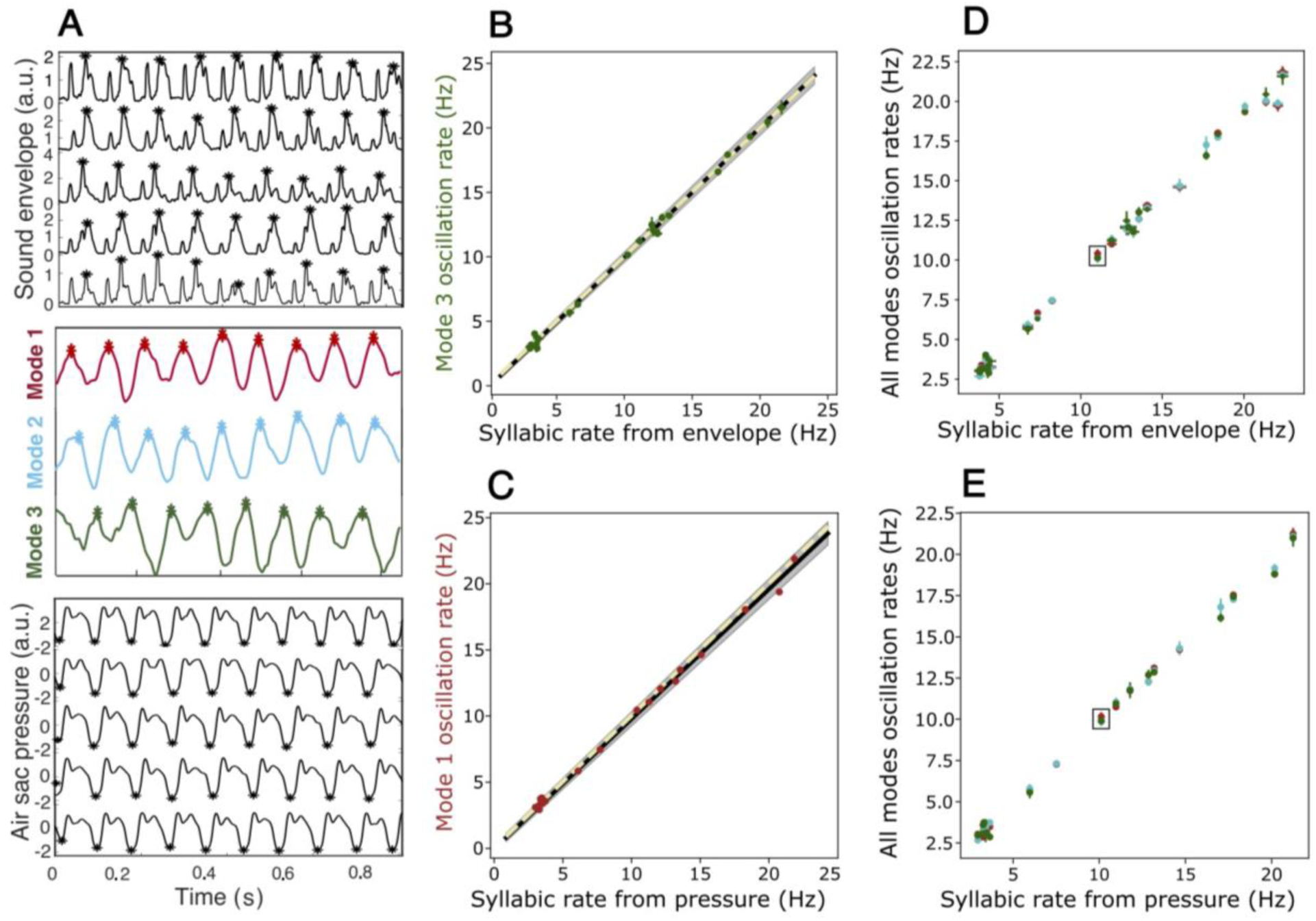
Oscillation frequency of modes matches syllabic rate. **A)** The (Q) phrase of caFLL001-VioAma is shown as an example, with asterisks indicating the detected events used for calculating the frequencies adjusted in the linear regressions. *Top panel:* 5 examples of the sound envelope. *Middle panel:* modes obtained with the autoencoder network. *Bottom panel:* 5 examples of the pressure recording. **B and C)** Linear regression between the oscillation rate of mode 3 vs the syllabic rate calculated from the envelope (panel B) and between the oscillation rate of mode 1 vs the syllabic rate calculated from the air sac pressure (panel C). The points with error bars represent the frequencies with their standard error. The black line represents the fit, the gray shading the 95% confidence interval, and the yellow dashed line is the identity line (y=x). The identity line is significantly contained within the confidence interval of the fit in both cases (p>0.9, ad-hoc test, supplementary figure S10). Regression parameters: in B) slope=1.00 (95% CI =[0.99-1.02]), intercept=0.07 (95% CI=[-0.09-0.13]), R^2^=0.9999 (24 total points, 4 birds); in C) slope=0.98 (95% CI =[0.96-1.00]), intercept=-0.13 (95% CI=[-0.32, 0.30]), R^2^=0.9968 (17 total points, 2 birds). **D and E)** Oscillation rates of all modes vs syllabic rate calculated from the sound envelope (panel D) or air sac pressure (panel E). The points overlap significantly, forming a line, and the frequency distributions in each mode show no significant differences (Kruskal-Wallis test, p-value>0.5, supplementary figure S12). The points corresponding to the example shown in panel A are highlighted with a box. The points with error bars represent the frequencies with their standard error. Red: mode 1, Light blue: mode 2, Green: mode 3. These results show that the frequency at which the modes oscillate closely matches the song syllabic rate.

Linear regression analyses comparing syllabic rates derived from the sound envelope with the oscillation frequencies of the latent modes are shown in Figure 4B and Supplementary Figures S10A and S10C. Agreement was evaluated by testing whether the relationship between both variables approached the identity function. Because both syllabic rate (behavioral measure) and mode frequency (latent representation) are subject to measurement error, orthogonal regression was used. Given the limited number of data points per regression (< 30), confidence intervals were estimated using residual bootstrapping (see Methods). The dataset spans a broad frequency range (2 - 27 Hz), highlighting the diversity in the rhythmic behavior. Standard diagnostic tests did not reject normality or homoscedasticity assumptions, as confirmed by histograms, Q–Q plots, and residual-versus-predicted value plots (Supplementary Figure S11).

Figure 4B shows the regression between syllabic rate (sound envelope-derived) and oscillation frequency for one representative mode (N = 24, 4 birds), yielding a slope of 1.00 (95% CI: [0.99, 1.02]), intercept of 0.07 Hz (95% CI: [–0.09, 0.13] Hz), and R² = 0.9999. Similar results were obtained for the other two modes (slopes ≈ 0.99, intercepts near 0, R² > 0.999; Supplementary Figures S10A and S10C), indicating a linear relationship across the 2–27 Hz range. To formally test whether this relationship was statistically indistinguishable from the identity line (slope = 1, intercept = 0), we performed a bootstrap-based analysis allowing a ±0.05 margin for both parameters. In all three modes, the identity line fell within the bootstrapped confidence intervals (p > 0.9; Supplementary Figure S10B–E). These results indicate that the oscillation frequencies of the latent modes closely match the syllabic rate of the song. Because the modes integrate activity across multiple stereotaxic coordinates of HVC, this finding supports the interpretation that rhythmic features of vocal behavior are encoded at the population level. Consistent with this interpretation, oscillation frequencies did not differ significantly across modes within the same phrase (Kruskal–Wallis test, p > 0.5; Supplementary Figure S12), indicating that all three modes convey the same rhythmic information. This overlap is evident in Figures 4D and 4E.

Similar results were obtained when syllabic rates were derived from air sac pressure recordings (2 birds; Figure 4C and Supplementary Figure S10F–H). Slopes ranged from 0.95 to 0.98 with R² > 0.99. The identity line was contained within the confidence intervals for modes 1 and 2, whereas for mode 3 the intercept slightly deviated beyond the ±0.05 criterion (p << 0.01), although the slope remained within range. As in the envelope-based regressions, we evaluated whether the identity line (slope = 1, intercept = 0) fell within the bootstrapped confidence intervals. This criterion was satisfied for modes 1 and 2 (Supplementary Figures S10G and S10J). For mode 3, however, the intercept consistently fell outside the ±0.05 margin (p << 0.01; Supplementary Figure S10I, lower panel), whereas the slope remained within range (upper panel).

We next compared the latent modes with a more conventional approach based on averaging all MUA traces, a common strategy to increase signal-to-noise ratio. After z-score normalization (mean subtraction and division by standard deviation), we quantified local oscillation amplitude for the mean trace and the three modes. A one-way ANOVA revealed significant differences across traces (p < 0.05). Post hoc Tukey tests showed that the averaged MUA trace had significantly lower local amplitude than each of the three modes, whereas no significant differences were observed among the modes themselves. Mean local amplitudes were: Xmean = (1.41 ± 0.24)𝛔, Xmode1 = (3.04 ± 0.24)𝛔, Xmode2 = (2.88 ± 0.24)𝛔, and Xmode3 = (2.77 ± 0.24)𝛔. These results indicate that dimensionality reduction preserves and enhances oscillatory structure that is partially attenuated by simple averaging.

Together, these findings show that autoencoder-derived latent modes offer a compact, low-dimensional representation of neural population dynamics that codes for the rhythmic structure of the vocal behavior.

## Conclusions

Understanding how neural activity encodes behavior along the song system has long been a central challenge in neuroscience. Our results demonstrate that neural population activity in HVC during singing can be captured within a compact low-dimensional space that preserves key features of vocal behavior. Using an unsupervised autoencoder applied to multi-unit recordings, we identified latent modes whose oscillation frequencies closely match both the syllabic repetition rate of the song and the underlying respiratory motor pattern.

Importantly, our results in canaries show multiunit activity in HVC already reflecting salient rhythmic features of song, indicating that population activity contains a rich representation of behaviorally relevant motor structure. These findings extend previous observations obtained during passive playback to the actively behaving individual, showing that structured rhythmic dynamics are robustly present during natural song production.

More broadly, this work supports a view in which complex motor behaviors emerge from low-dimensional neural dynamics, where a small number of collective variables capture essential features of high-dimensional population activity. Within this framework, rhythmic behaviors such as birdsong may be understood as trajectories constrained by dynamical structure rather than by sequential activation alone.

Finally, the approach introduced here provides a general and data-driven strategy to extract interpretable dynamical structure from large-scale neural recordings. By combining dimensionality reduction with behaviorally grounded validation, this framework opens the door to comparative studies across species and motor systems, and offers a promising avenue for investigating how neural populations generate and control temporally structured behaviors.

## Methods Subjects

The data were collected from four adult male canaries (*Serinus canaria*) obtained from a local supplier (caCTH2447-RoNa, caCTH03-VioAma, caCTH140-VioVe, caFLL001-VioAma). All data used in this manuscript were acquired between late October and early March—a period during which canaries perform their mating season songs in the Southern Hemisphere. Birds were individually housed throughout the experiment, fed *ad libitum*, and kept on a 14:10-hour light-dark cycle matching the summer daylight cycle in Argentina. Experiments were conducted under a protocol approved by the Institutional Animal Care and Use Committee (C.I.C.U.A.L.) of the University of Buenos Aires, Argentina (FCEN-UBA).

## Experimental Methods

### Neuronal Activity

Extracellular neuronal activity in the telencephalic nucleus HVC (proper name) and sound were recorded simultaneously during song production, as described by Herbert et al. (2020). To promote singing, birds were intermittently exposed to visual and auditory stimuli from other canaries. Recordings were performed for as long as the subjects exhibited high-quality physiological signals and frequent singing. Recording durations were 21 days for three birds and 48 days for the fourth.

Neuronal activity, sound, and air sac pressure were continuously monitored, with recordings triggered by sustained sound above a manually calibrated decibel threshold, corresponding to singing. Electrophysiological recordings of neuronal populations were made using low-impedance microwire tetrode arrays fabricated in-house from 12.7 μm diameter polyimide-coated tungsten wires (Tungsten 99.95% CS Wire, California Fine Wire) [Henze et al., 2000].

The tetrodes were mounted on a lightweight manual microdrive adapted from [Vandecasteele et al., 2012]. Differential recordings were obtained between tetrode wires and a reference electrode implanted outside the HVC, in addition to a silver ground wire placed between the skull’s lower layer and the dura mater. The recording device measured 1 cm in height, weighed approximately 0.53g, and included an additional 0.79g headstage. Neural signals were amplified on an RHD2132 16-channel amplifier board, and song, neural, and pressure data were recorded at 30 kS/s using an Intan Technologies RHD2000 USB interface board.

### MUA Traces

To extract patterns of neuronal population spiking activity during singing, the recorded extracellular voltage was digitally filtered with a high-pass filter (300 Hz cutoff frequency, third-order Butterworth IIR filter). Disconnected or noisy channels were discarded by visual inspection, checking impedances and voltage range (very high impedances and amplitudes correspond to such channels). This neuronal signal was filtered with a threshold thr = -3σ = -3|x|/0.6745, where σ is a robust estimator for the background noise standard deviation (Quiroga *et al*., 2004), and x is the high-pass filtered data. This threshold captured most spikes without introducing significant noise that would interfere with the signal pattern. This relative amplitude threshold was implemented to detect multi-unit activity (MUA) peaks, which exhibited different spike shapes, amplitudes, and presences across the channels of each tetrode. MUA traces were computed by smoothing the spiking data using a kernel density estimation (Gaussian kernel with a bandwidth of 5 ms, corresponding to 150 samples at a 30 kHz sampling rate, evaluated at the original temporal resolution of the data, 0.033 ms). This procedure yielded continuous MUA traces sampled at 30 kHz. The resulting smoothed trace was multiplied by the spike count to obtain an absolute measure of spiking rate. Parameters for this smoothing were determined by visual inspection to preserve temporal patterns of neuronal activity (periods of activity-silence) while minimizing noise across all phrase types, considering their different temporal scales (2 to 27 Hz). The resulting smoothed trace was multiplied by the spike count to obtain an absolute measure of spiking rate. We obtained one MUA trace per channel for each rendition of the song segments (16 MUA traces in total). MUA traces corresponding to each tetrode were averaged to obtain 4 traces per song rendition. All analyses were performed offline in Matlab (MathWorks, www.mathworks.com).

### Song

Audio was acquired using a 20Hz - 20kHz electret microphone mounted on a Maxim MAX 4466 amplifier board with adjustable gain placed inside the sound attenuating chamber where the bird cage was placed. The audio signal was recorded on an analog input of the RHD2000 USB interface board and was sampled at the same rate as the neural signal. The audio signal was digitally high-pass filtered (300 Hz cutoff frequency, third order Butterworth) in Matlab. Spectrograms of the audio signal were computed using Matlab with a 10ms gaussian window and temporal overlap of 95%.

### Air Sac Pressure

Air sac pressure was recorded using a flexible cannula and a miniature piezoresistive pressure transducer (Fujikura model FPM-02PG). For further details see for example (Alliende et al. 2010). Pressure data was amplified with custom hardware and acquired into an analog input of the RHD2000 USB interface board with the same rate as the neural signal. The pressure signal was digitally low-pass filtered (300Hz cutoff frequency, third order Butterworth filter) in Matlab.

## Data Analysis

### Preparation of data to be entered into the autoencoder network

All phrases used for analysis were identified with a "label" referring to the individual repertoire of each bird, cataloged according to the acoustic and rhythmic characteristics of the individual’s song. Phrases from each bird’s repertoire that were sung in all recording sites (different depths within the HVC) were selected. For each syllable identity, the minimum duration of the phrase across all sites was used to determine the maximum length of the segments (time-windows cropped from phrases) used for analysis. The cap was 1 s for the longest phrases. All segments were aligned at the first syllable of the phrase and extended to the target duration. Segments of different syllable identities were concatenated in the same order for each site before analysis with the autoencoder.

## Autoencoder Network

The autoencoder was implemented in Python and consisted of fully connected layers with ReLU activation functions, except for the linear units in the middle layer. The network was trained to minimize the mean squared error (MSE) using the Adam optimizer (batch size = 512, 50 epochs). Training was stopped when the average percentage change in test MSE fell below 1%. The dimensionality of the input layer corresponded to the number of valid recording sites per bird, defined as the total number of sites recorded minus discarded tetrodes. The total number of recorded sites was given by the number of explored depths multiplied by four (the number of tetrodes per depth). Accordingly, the input dimensionality was 44 for caCTH03-VioAma (each trace comprising 142505 samples), 48 for caCTH2447-RoNa (142207 samples), 29 for caCTH140-VioVe (93905 samples), and 52 for caFLL001-VioAma (331814 samples).

## Processing of audio recordings

### Audio Recordings

Sound envelopes were calculated using Matlab’s envelope function with the RMS method, chosen for its robustness in capturing oscillation patterns while minimizing noise. The sliding window sizes were adjusted based on the syllable oscillation frequencies: 200 points for the fastest oscillations (8 to 27 Hz) and 500 points for the slowest (2 to 8 Hz). Additional smoothing was applied to ensure accurate peak detection in slower oscillation signals.

## Processing of autoencoder latent modes

### Mode traces

In order to identify peaks, mode traces were segmented by syllable type and smoothed using Matlab’s smoothdata function with a sliding mean, this helped avoid detecting noise-related peaks in the signal. Window lengths were adjusted proportionally to syllable oscillation periods to optimize peak detection, maintaining representative peaks, and reducing noise in slower oscillation signals.

### Detection of syllable onset in oscillations

#### a) Modes and sound envelope

Onset detection was based on abrupt changes in oscillation patterns. Local maxima and minima were detected independently with Matlab’s findpeaks function, ensuring that they were spaced by at least the duration of the syllable (which was determined manually through visual inspection). The differences between the detected maxima and minima were then computed. These differences represent the changes in oscillation between successive peaks. After calculating these differences, they were ordered from highest to lowest to retain only the most significant changes. This allowed us to focus on the most prominent events, reducing the possibility of false detections due to smaller, less meaningful oscillations.

Signals classified as noisy were excluded from the analysis. A noisy signal was defined as one that exhibited irregular oscillations, with more than four peaks surpassing the median of the FFT transformation.

Periods were calculated for all phrases in sound envelopes corresponding to the recorded depths (caCTH2447-RoNa = 13, caCTH03-VioAma = 10, caCTH140-VioVe = 9, caFLL001-VioAma = 13; see supplementary figures S1 to S4 for detailed envelopes). The period was defined as the time difference between the onset of one syllable and the next. These periods evolved randomly throughout the course of the phrases, and the mean and standard deviation of all the periods within each phrase were calculated to provide a comprehensive measure of the variation in syllable timing.

#### Pressure signal

Local minima (corresponding to inspiration) were detected. A sample of pressure signals equivalent in size to the number of analyzed sound envelopes was taken (10 samples, see supplementary figures S5 and S6). The individual periods were calculated as the difference between one inspiration and the next one.

## Signal Amplitude Quantification

The amplitudes of the traces were normalized using z-score normalization to express values in terms of standard deviations. The local amplitude of each oscillation was calculated as the average difference between the standard deviation values of peaks and valleys along the trace. These values were then grouped by phrase.

## Statistical Analyses

### Mode Oscillation Frequency vs. Syllabic Rate

Individual orthogonal regressions were performed for each mode using the ODR function from Python’s Scipy library:

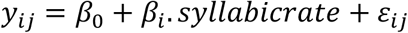

where 𝑦_𝑖𝑗_ represents the frequency of oscillations of the modes in the group i (each phrase); 𝛽_0_ is the intercept, representing the expected oscillation frequency when the syllabic rate is zero (non-interpretable in this context since the measurement range of frequencies was between 2 Hz and 25 Hz and 0 has no biological sense); 𝛽_𝑖_ is the coefficient representing the relationship between the mode frequency and the oscillation of envelope/air sac pressure frequency; 𝜀_𝑖𝑗_ is the residual error, which follows a normal distribution 𝜀_𝑖𝑗_ ∼ NID(0, σ²).

Standard errors were used as weights, and given the low sample size (#points_mode1/envelope_= 26, #_pointsmode2/envelope_= 27, #points_mode3/envelope_= 24; #points_mode1/pressure_= 17, #points_mode2/pressure_= 18, #points_mode3/pressure_= 16), bootstrapping was employed to obtain robust confidence intervals, repeating the process 1,000 times.

Normality and homoscedasticity assumptions were evaluated with statsmodels and scipy.stats, averaging p-values across the 1,000 samples. On average, the hypotheses of normality (Shapiro-Wilk, Anderson-Darling) and homoscedasticity (Breusch-Pagan, White) were not rejected (p-value > 0.05 in all cases). Results were visually corroborated using histograms, Q-Q plots, and residuals vs. predicted value plots.

To evaluate the inclusion of the identity line in linear fits, an ad hoc test was performed considering m=1±0.05 and b=0±0.05. This margin of error accounted for variability in singing behavior.

Finally, oscillation frequencies across the three modes were compared using the Kruskal-Wallis test, as some distributions did not meet normality (Shapiro-Wilk, p-value > 0.05), although variances were homogeneous (Levene, p-value > 0.05).

### Local Amplitude Comparison via ANOVA

A one-way ANOVA was performed to compare local amplitudes across four levels (average, mode 1, mode 2, and mode 3):

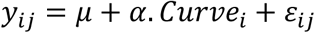

where 𝑦_𝑖𝑗_ represents the mean local amplitude for each phrase; 𝜇: General mean; .𝛼. 𝐶𝑢𝑟𝑣𝑒_𝑖_: Fixed effect of trace type (i=average, mode 1, mode 2, mode 3); 𝜀_𝑖𝑗_: Random error ((𝜀_𝑖𝑗_ ∼ *NID*(0, σ))

Normality and homoscedasticity assumptions were verified, and no evidence to reject them was found. Post-hoc contrasts were performed using Tukey’s method.

## Acknowledgments

The work was partially funded by Universidad de Buenos Aires (UBACYT-2023, 20020220300061BA), Agencia Nacional de Promoción de la investigación, el Desarrollo Tecnológico y la Innovación (PICT-2021-I-A-00965), and CONICET (KE3-11220210100475CO) (Argentina).

---

**Supplementary Figure S1.**
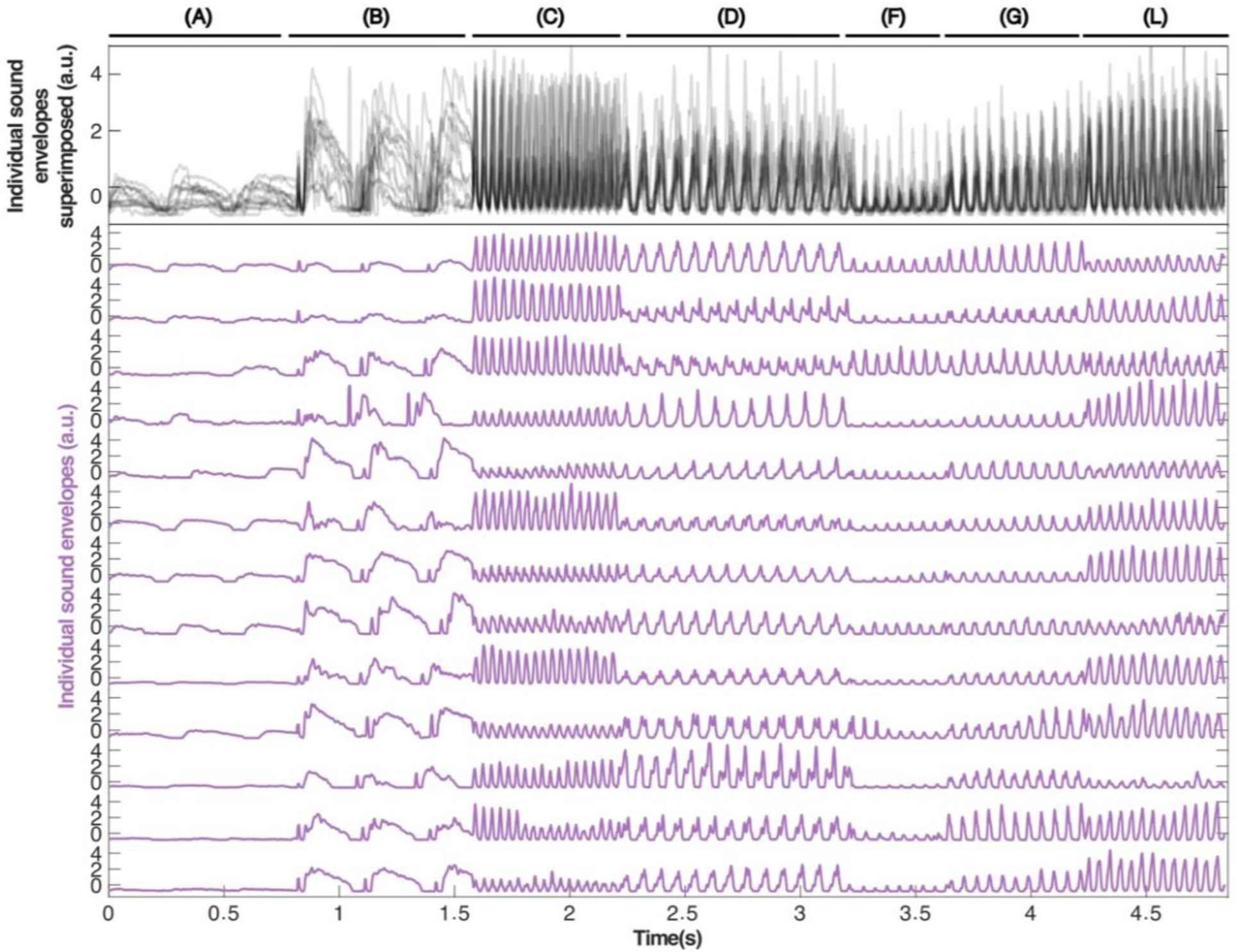
Individual sound envelopes, bird caCTH2447-RoNa. The traces shown in this figure correspond to the calculation of the sound envelope for each song performed by the bird during the neuronal activity recordings taken at different stereotaxic coordinates (for more details, see methods section). The classification of the phrases in the bird’s repertoire is shown at the top of the figure with a line and the corresponding labelling (an identifier with a letter). **Top panel:** all sound envelopes superimposed (black lines with transparency), with darker areas corresponding to moments of greater trace overlap, illustrating that the song is highly stereotyped, although with slight differences due to small variations in each execution. **Bottom panel:** each sound envelope is shown individually (violet traces), with each row representing the sound envelope of the song performed during the neuronal recording at each analyzed depth. It can be seen that each phrase is sung with a characteristic rhythmicity (syllabic rate) and the syllables of the same phrase are very similar when sung at different times (for example, the syllables of phrase D all have two notes, reflected in the envelope as a double peak). Songs performed on different days have similarities that justify grouping them into categories (phrase) but variations such that individual event detection is needed in each trace to calculate the syllabic rate.

**Supplementary Figure S2.**
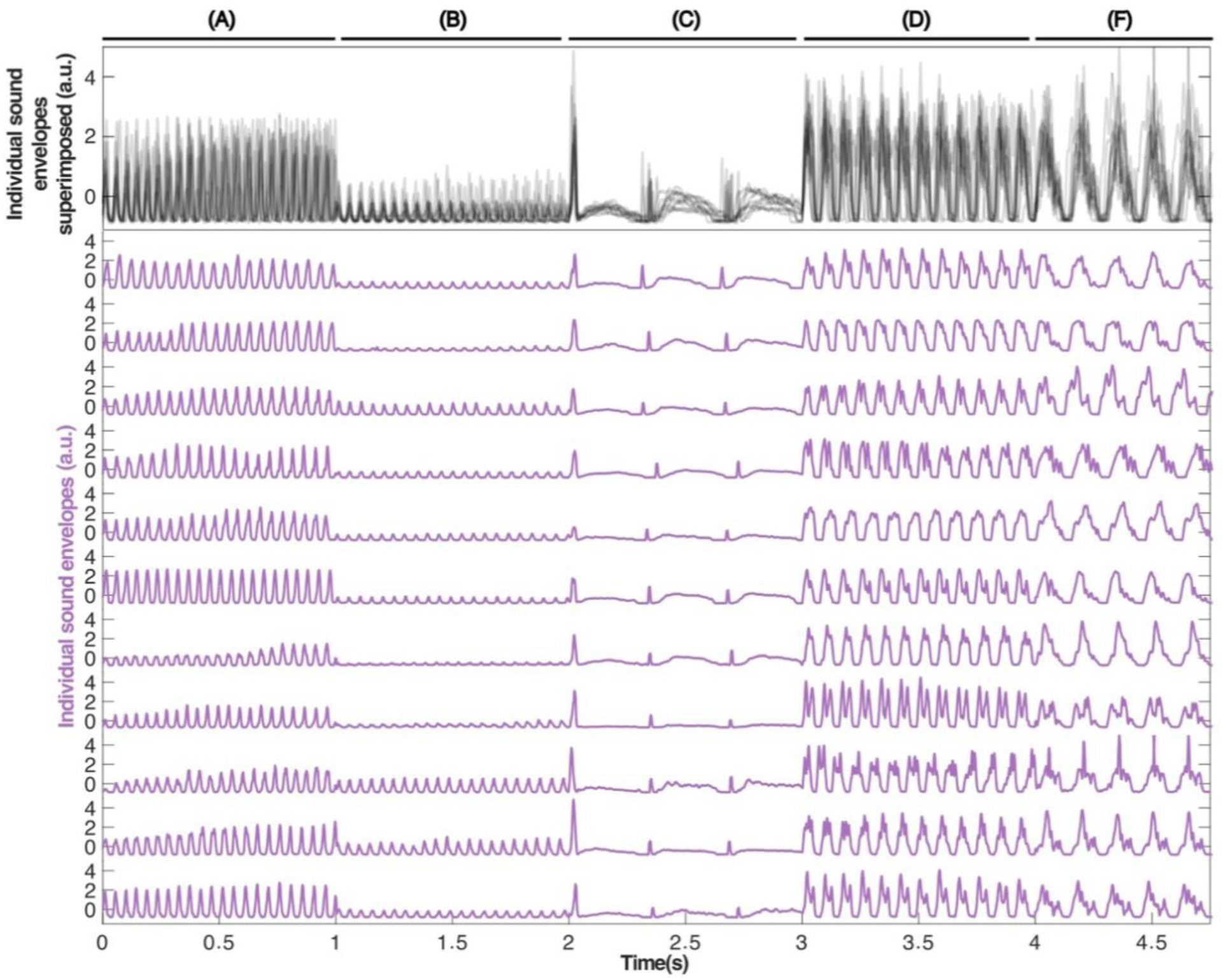
Individual sound envelopes, bird caCTH03-VioAma. The traces shown in this figure correspond to the calculation of the sound envelope for each song performed by the bird during the neuronal activity recordings taken at different stereotaxic coordinates (for more details, see methods section). The classification of the phrases in the bird’s repertoire is shown at the top of the figure with a line and the corresponding labelling (an identifier with a letter). **Top panel:** all sound envelopes superimposed (black lines with transparency), with darker areas corresponding to moments of greater trace overlap, illustrating that the song is highly stereotyped, although with slight differences due to small variations in each execution. **Bottom panel:** each sound envelope is shown individually (violet traces), with each row representing the sound envelope of the song performed during the neuronal recording at each analysed depth. It can be seen that each phrase is sung with a characteristic rhythmicity (syllabic rate) and the syllables of the same phrase are very similar when sung at different times (for example, the syllables of phrase F all have two notes, reflected in the envelope as a double peak). Songs performed on different days have similarities that justify grouping them into categories (phrase) but variations such that individual event detection is needed in each trace to calculate the syllabic rate.

**Supplementary Figure S3.**
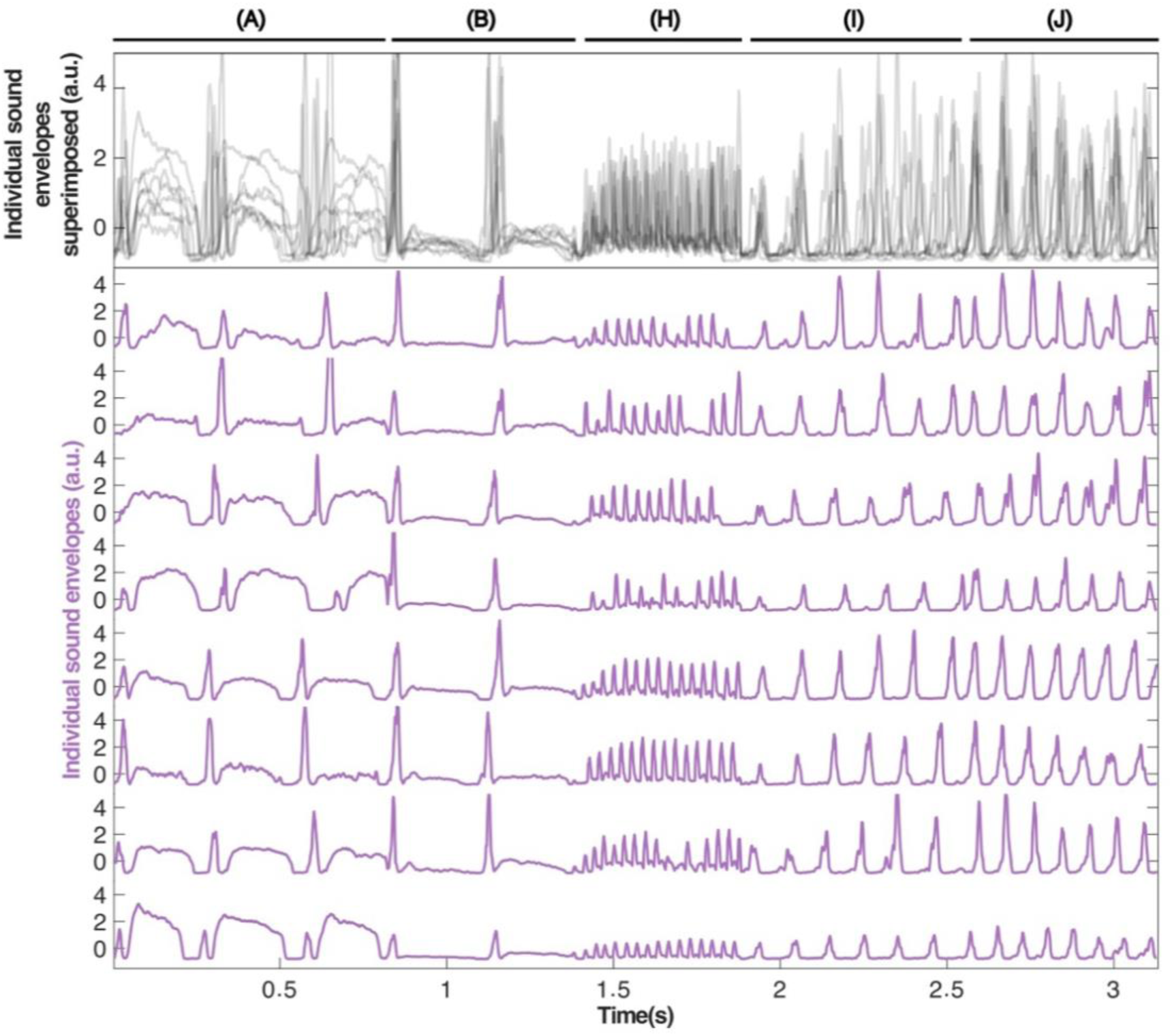
Individual sound envelopes, bird caCTH140-VioVe. The traces shown in this figure correspond to the calculation of the sound envelope for each song performed by the bird during the neuronal activity recordings taken at different stereotaxic coordinates (for more details, see methods section). The classification of the phrases in the bird’s repertoire is shown at the top of the figure with a line and the corresponding labelling (an identifier with a letter). **Top panel:** all sound envelopes superimposed (black lines with transparency), with darker areas corresponding to moments of greater trace overlap, illustrating that the song is highly stereotyped, although with slight differences due to small variations in each execution. **Bottom panel:** each sound envelope is shown individually (violet traces), with each row representing the sound envelope of the song performed during the neuronal recording at each analysed depth. It can be seen that each phrase is sung with a characteristic rhythmicity (syllabic rate) and the syllables of the same phrase are very similar when sung at different times (for example, the syllables of the phrases A and B have two notes, reflected in the envelope as a small peak at the beginning and a wider trace later). Songs performed on different days have similarities that justify grouping them into categories (phrase) but variations such that individual event detection is needed in each trace to calculate the syllabic rate.

**Supplementary Figure S4.**
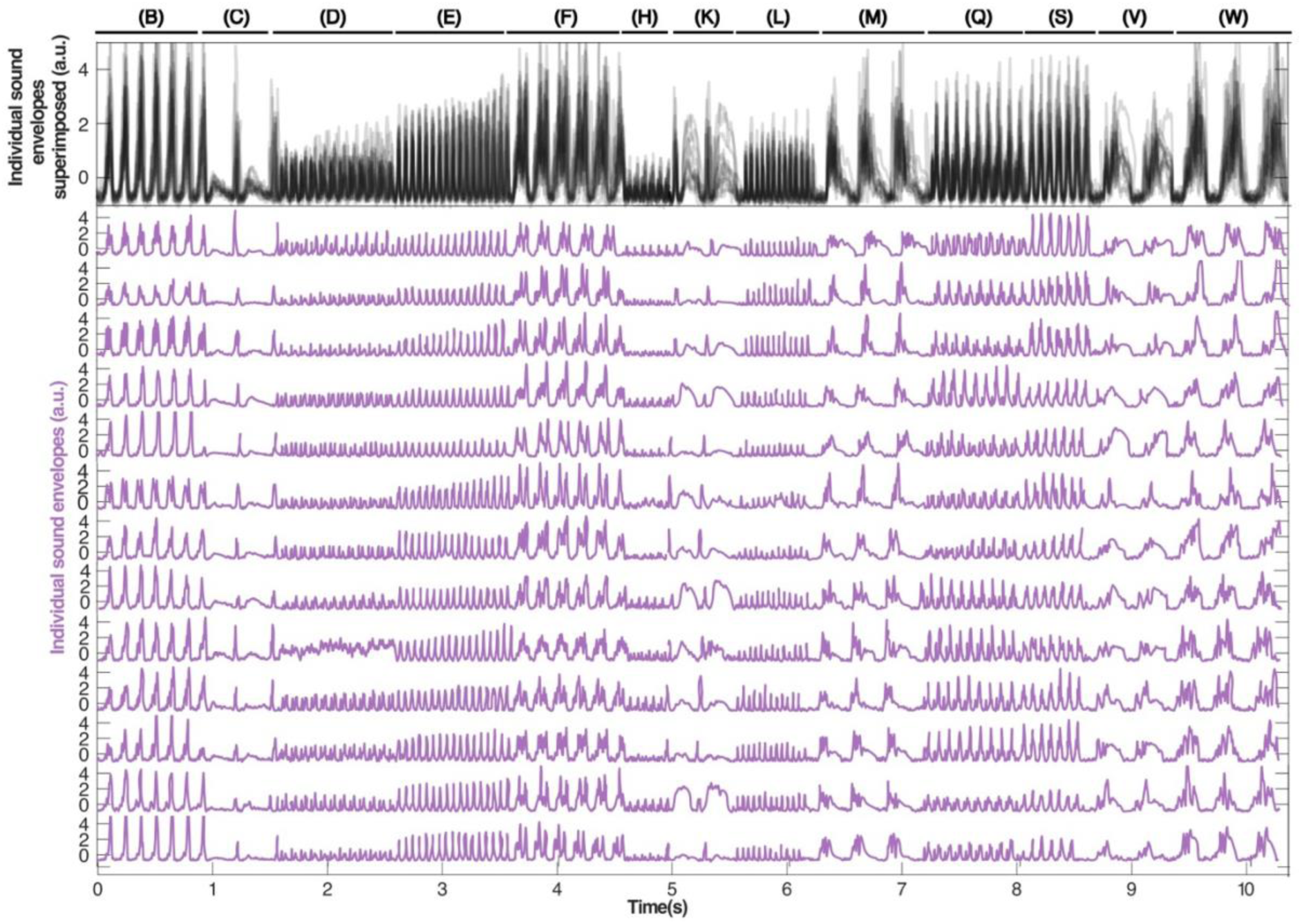
Individual sound envelopes, bird caFLL001-VioAma. The traces shown in this figure correspond to the calculation of the sound envelope for each song performed by the bird during the neuronal activity recordings taken at different stereotaxic coordinates (for more details, see methods section). The classification of the phrases in the bird’s repertoire is shown at the top of the figure with a line and the corresponding labelling (an identifier with a letter). **Top panel:** all sound envelopes superimposed (black lines with transparency), with darker areas corresponding to moments of greater trace overlap, illustrating that the song is highly stereotyped, although with slight differences due to small variations in each execution. **Bottom panel:** each sound envelope is shown individually (violet traces), with each row representing the sound envelope of the song performed during the neuronal recording at each analysed depth. It can be seen that each phrase is sung with a characteristic rhythmicity (syllabic rate) and the syllables of the same phrase are very similar when sung at different times (for example, the three peaks of phrase F or the narrow peak followed by a wider peak of phrase K). Songs performed on different days have similarities that justify grouping them into categories (phrase) but variations such that individual event detection is needed in each trace to calculate the syllabic rate.

**Supplementary Figure S5.**
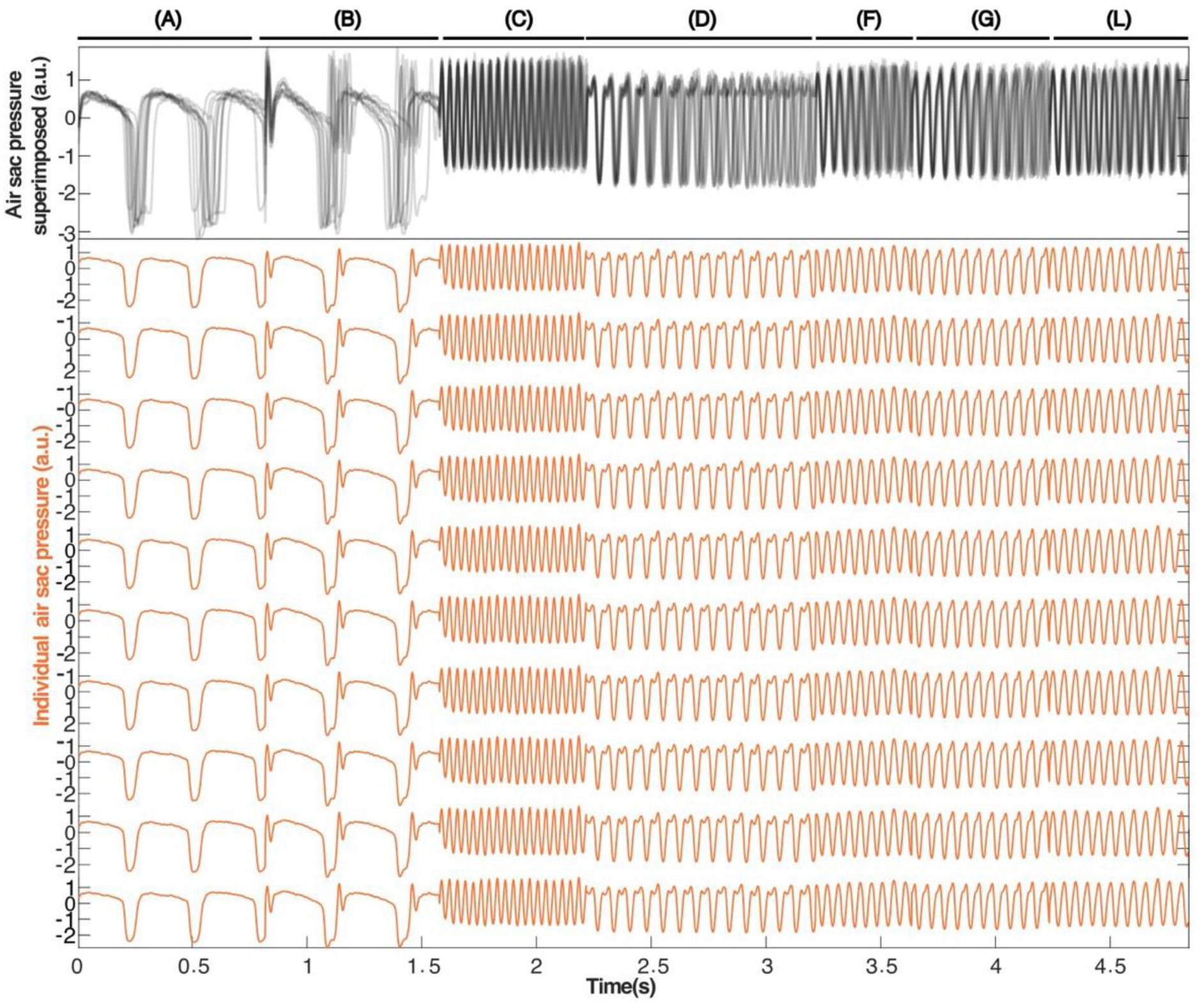
Pressure recordings, bird caCTH2447-RoNa. The traces shown in this figure correspond to the air sac pressure recordings used to calculate the syllabic rate from the pressure, performed on days following the neuronal recordings (N=10 samples of each phrase). The classification of the phrases in the bird’s repertoire is shown at the top of the figure with a line and the corresponding labelling (an identifier with a letter). **Top panel:** all traces superimposed (black lines with transparency), with darker areas corresponding to moments of greater overlap, illustrating that the motor gesture underlying the song is highly stereotyped. **Bottom panel:** each pressure recording is shown individually (orange traces), with each row representing an example constructed by concatenating the pressure recording corresponding to the different phrases used as input to the autoencoder network. It can be seen that each phrase is sung with a characteristic rhythmicity (syllabic rate) and the syllables of the same phrase are very similar when sung at different times. Songs performed on different days have similarities that justify grouping them into categories (phrase) but variations such that individual event detection is needed in each trace to calculate the syllabic rate.

**Supplementary Figure S6.**
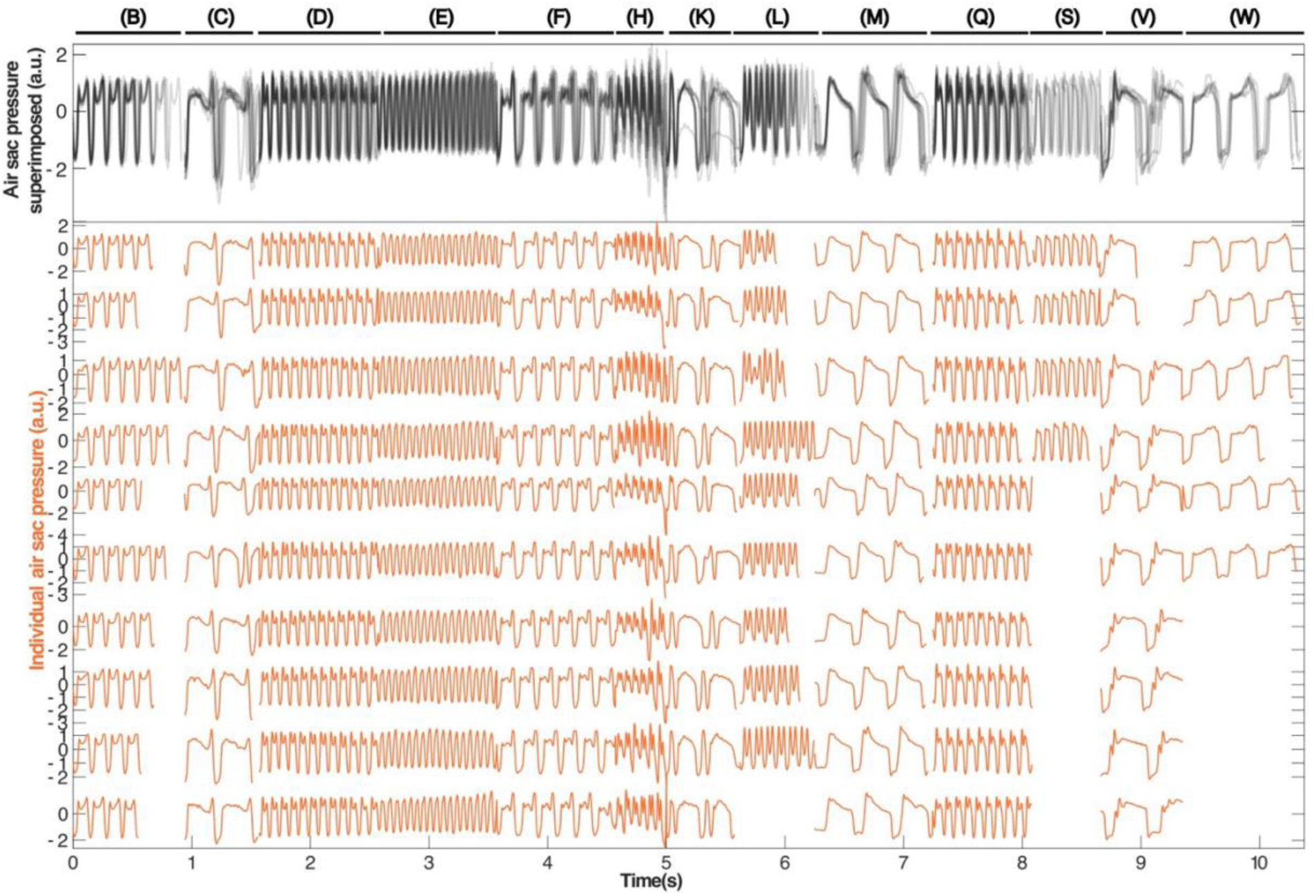
Pressure recordings, bird caFLL001-VioAma. The traces shown in this figure correspond to the air sac pressure recordings used to calculate the syllabic rate from the pressure, performed on days following the neuronal recordings (N=10 samples of each phrase). Since there are not 10 examples of each phrase (some are sung with lower probability than others, and the days with pressure recordings were fewer), the figure shows some blank spaces in the phrases where the total number of examples sought could not be found. The classification of the phrases in the bird’s repertoire is shown at the top of the figure with a line and the corresponding labelling (an identifier with a letter). **Top panel:** all traces superimposed (black lines with transparency), with darker areas corresponding to moments of greater overlap, illustrating that the motor gesture underlying the song is highly stereotyped. **Bottom panel:** each pressure recording is shown individually (orange traces), with each row representing an example constructed by concatenating the pressure recording corresponding to the different phrases used as input to the autoencoder network. It can be seen that each phrase is sung with a characteristic rhythmicity (syllabic rate) and the syllables of the same phrase are very similar when sung at different times. Songs performed on different days have similarities that justify grouping them into categories (phrase) but variations such that individual event detection is needed in each trace to calculate the syllabic rate.

**Supplementary Figure S7.**
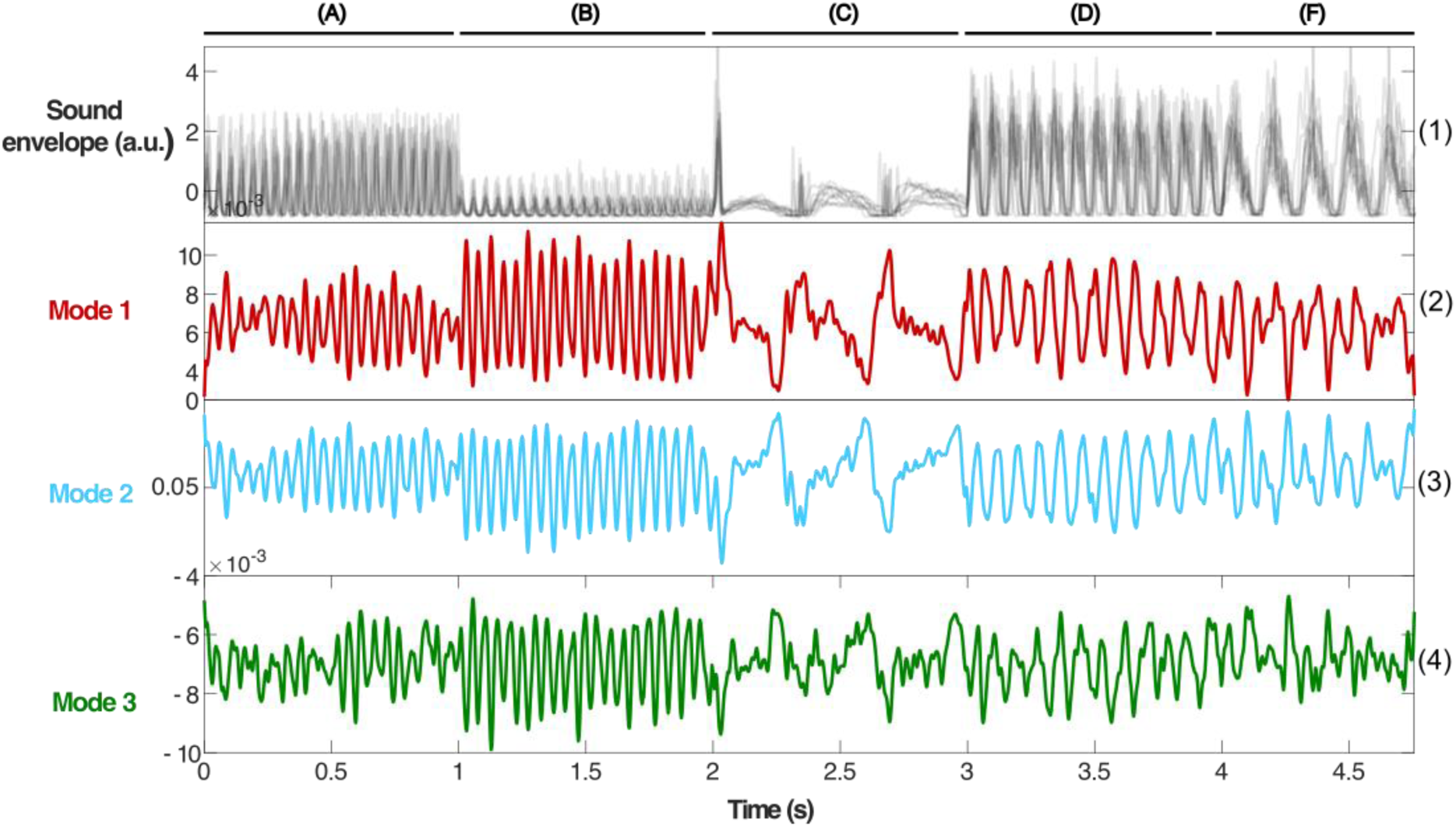
Autoencoder latent modes for bird caCTH03-VioAma. **Panel 1:** all normalized sound envelopes of all recorded stereotaxic coordinates used as input to the autoencoder network, superimposed with transparency. Darker areas correspond to moments where the traces overlap, showing that while there are slight rhythmic and amplitude variations, the song is highly stereotyped. The classification of the phrases in the bird’s repertoire is shown at the top of the figure with a line and the corresponding labelling (and an identifier with a letter). **Panels 2 to 4:** traces corresponding to each of the three modes of the network’s latent layer, the minimum information necessary to recover the input layer information with the least error. We can observe that, similar to individual MUA traces, each phrase has a characteristic oscillation frequency, and details of the sound envelope and/or air sac pressure structure are reflected in the modes.

**Supplementary Figure S8.**
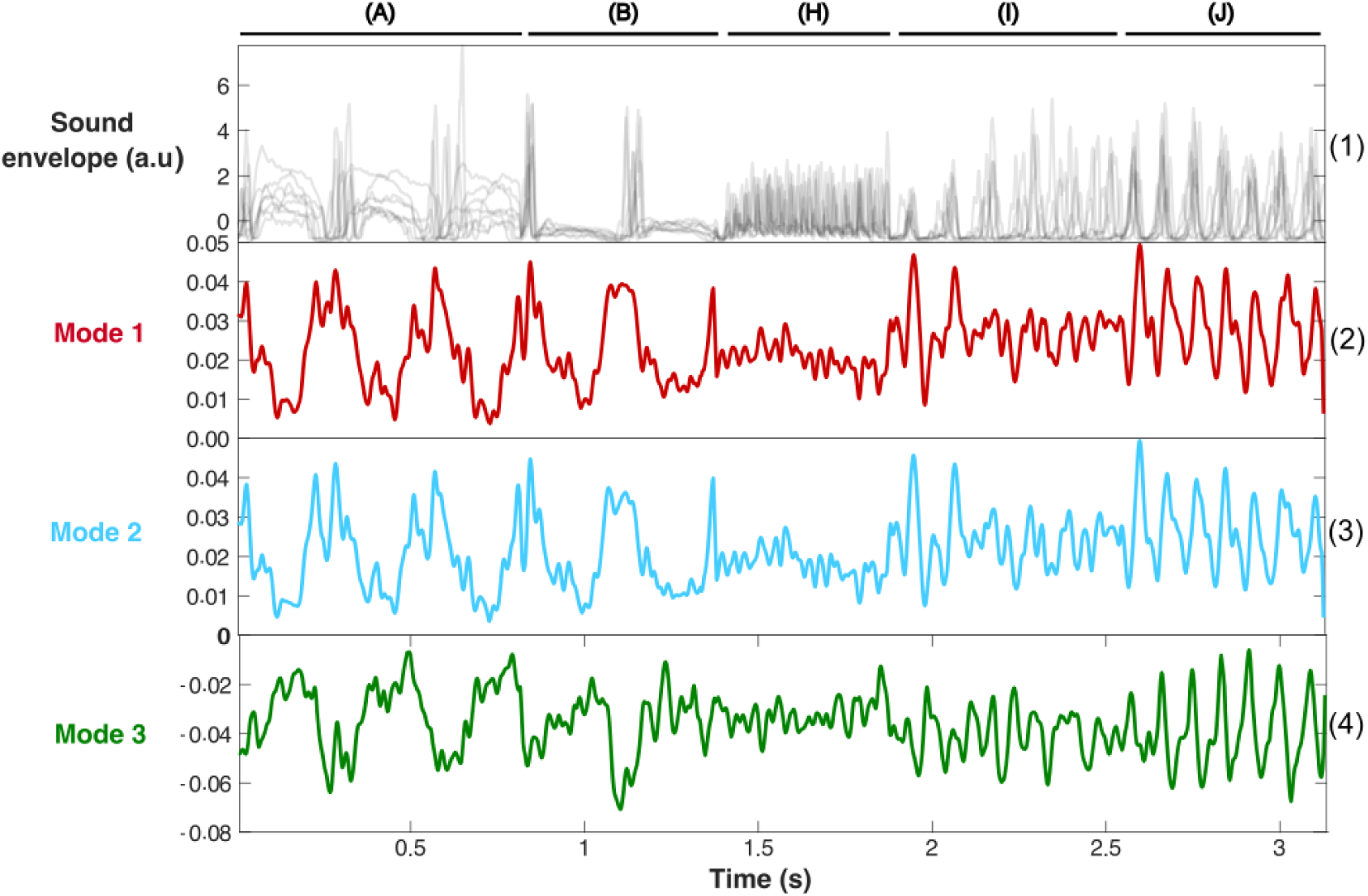
Autoencoder latent modes for bird caCTH140-VioVe. **Panel 1:** all normalized sound envelopes of all recorded stereotaxic coordinates used as input to the autoencoder network, superimposed with transparency. Darker areas correspond to moments where the traces overlap, showing that while there are slight rhythmic and amplitude variations, the song is highly stereotyped. The classification of the phrases in the bird’s repertoire is shown at the top of the figure with a line and the corresponding labelling (an identifier with a letter). **Panels 2 to 4:** traces corresponding to each of the three modes of the network’s latent layer, the minimum information necessary to recover the input layer information with the least error. We can observe that, similar to individual MUA traces, each phrase has a characteristic oscillation frequency, and details of the sound envelope and/or air sac pressure structure are reflected in the modes.

**Supplementary Figure S9.**
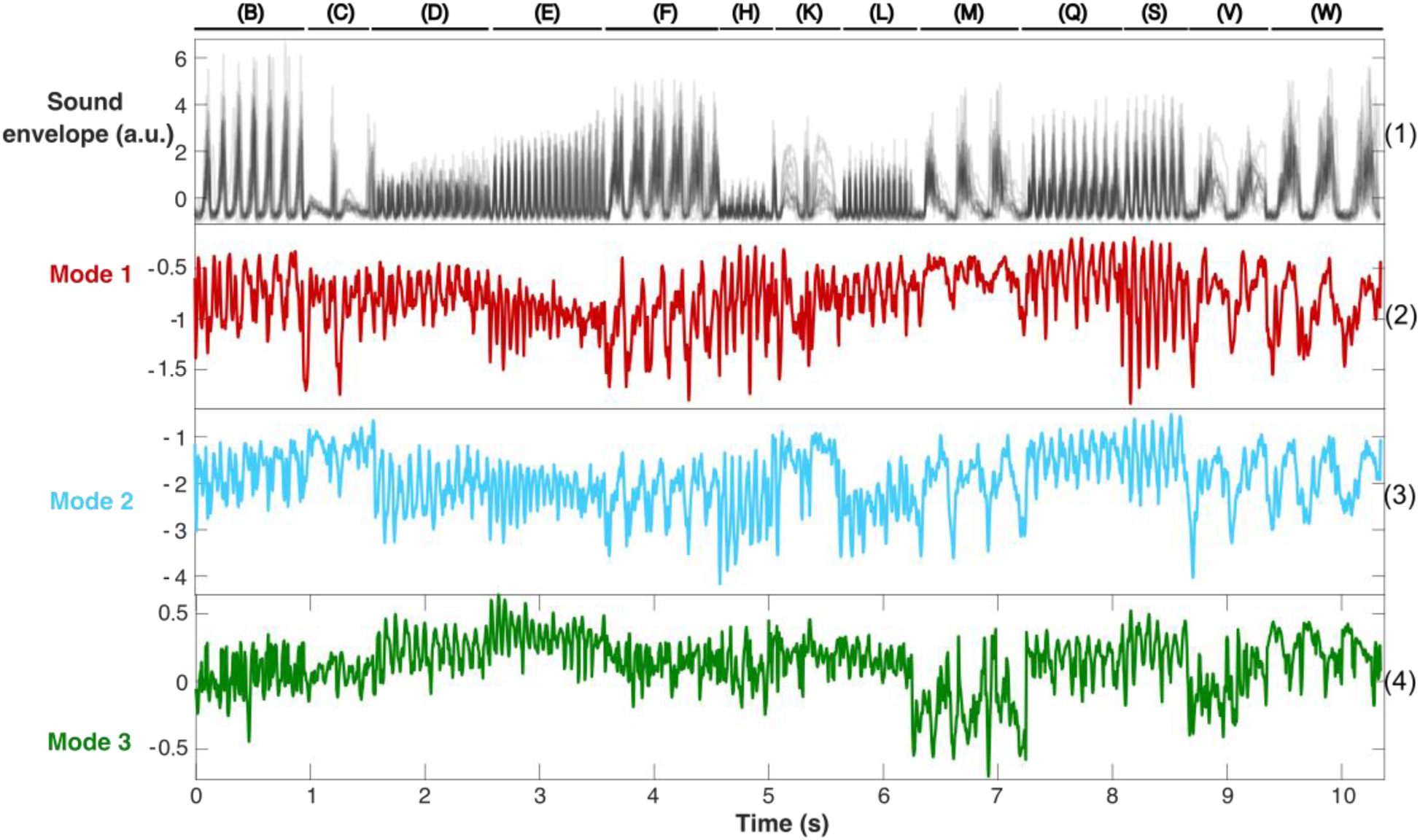
Autoencoder latent modes for bird caFLL001-VioAma. Panel 1: all normalized sound envelopes of all recorded stereotaxic coordinates used as input to the autoencoder network, superimposed with transparency. Darker areas correspond to moments where the traces overlap, showing that while there are slight rhythmic and amplitude variations, the song is highly stereotyped. The classification of the phrases in the bird’s repertoire is shown at the top of the figure with a line and the corresponding labelling (an identifier with a letter). **Panels 2 to 4:** traces corresponding to each of the three modes of the network’s latent layer, the minimum information necessary to recover the input layer information with the least error. We can observe that, similar to individual MUA traces, each phrase has a characteristic oscillation frequency, and details of the sound envelope and/or air sac pressure structure are reflected in the modes.

**Supplementary Figure S10.**
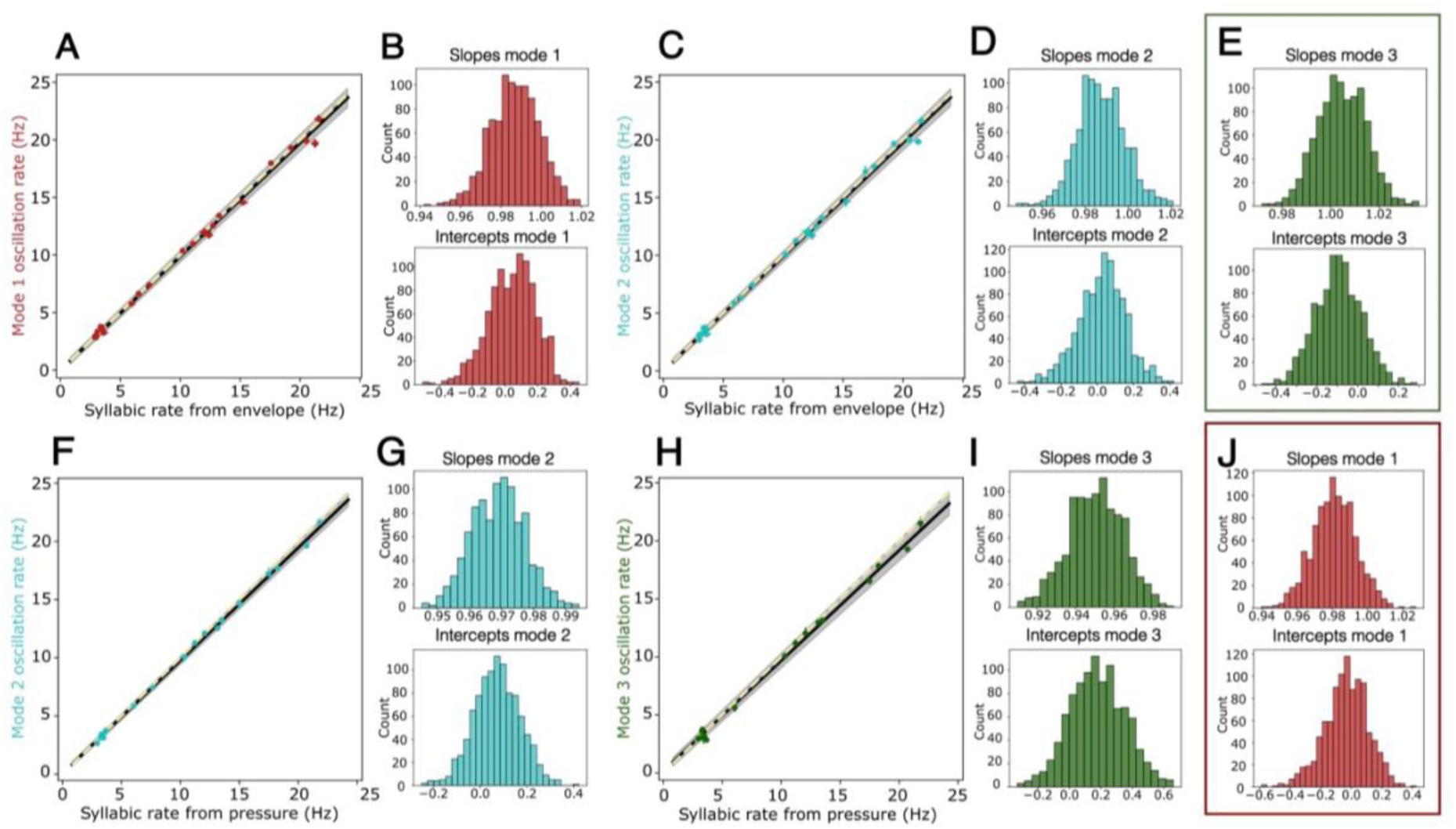
Oscillation frequency of modes vs syllabic rate. **A, C, F, and H)** Linear regression between the oscillation rate of the modes vs the syllabic rate calculated from the envelope (panels A and C) or from air sac pressure (panels F and H). Points with error bars represent the mean frequencies with their standard error. The black line represents the fit, the grey shading the 95% confidence interval, and the yellow dashed line is the identity line (y=x). **B, D, E, G, I, and J)** Distribution of regression parameters obtained in the 1000 iterations of the linear regression fit between mode frequencies vs syllabic rate calculated from the sound envelope (panels B, D, and E) or air sac pressure (panels G, I, and J) using bootstrapping of residuals. *Top panel:* slope distribution, the hypothesis that the value 1±0.05 is significantly contained in all cases could not be rejected (p-value>0.9, ad-hoc test). *Bottom panel:* intercept distribution, the hypothesis that the value 0±0.05 is significantly contained could not be rejected (p-value>0.9, ad-hoc test) except in the case of mode frequencies vs syllabic rate from pressure linear regression fit (panel I, bottom, p-value<0,01, ad-hoc test). The linear regressions corresponding to panels E and J are shown in figures 4B and 4D, respectively. Red: mode 1, Light blue: mode 2, Green: mode 3. The parameters of the linear functions shown in this figure are as follows: in A) slope=0.99 (95% CI =[0.96-1.01]), intercept=-0.15 (95% CI=[-0.25, 0.30]), R^2^=0.9998, (26 total points, 4 birds). In C) slope=0.99 (95% CI =[0.97-1.01]), intercept=0.07 (95% CI=[-0.26-0.27]), R^2^=0.9997 (27 total points, 4 birds). In F) slope=0.97 (95% CI =[0.95-0.99]), intercept=0.06 (95% CI=[-0.13-0.26]), R^2^=0.9973 (18 total points, 2 birds). In H) slope= 0.95 (95% CI =[0.92-0.98]), intercept=0.10 (95% CI=[-0.13- 0.51]), R^2^=0.9934 (16 total points, 2 birds).

**Supplementary Figure S11.**
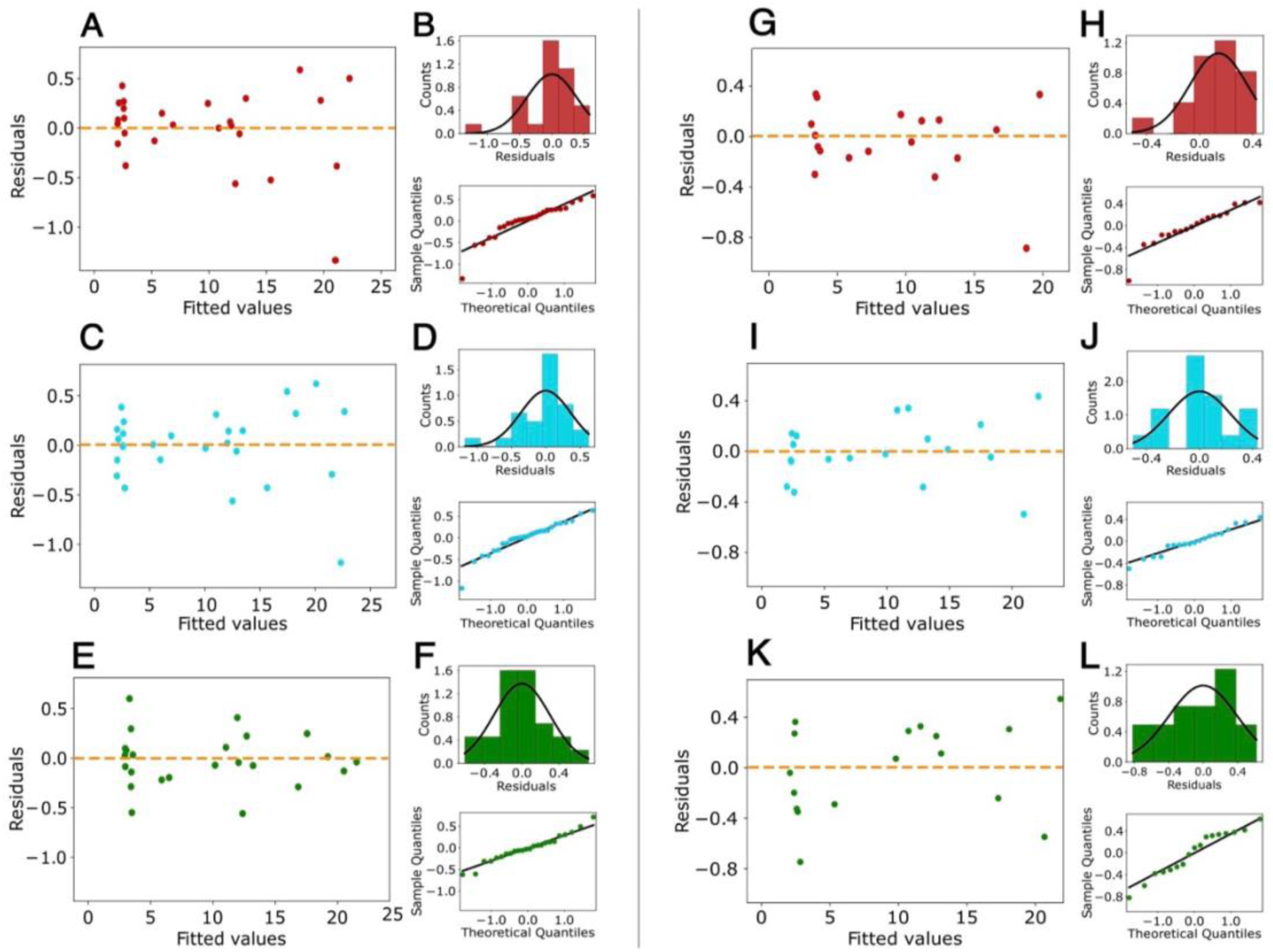
Goodness-of-fit tests for linear regressions between mode oscillation frequencies vs syllabic rate obtained from the envelope/pressure. **A, G, C, I, E, and K)** Residuals vs predicted distribution plot for the linear regression fit between mode frequencies and syllabic rates obtained from the envelope (A, C, and E) or air sac pressure (G, I, and K). Red: mode 1, Light blue: mode 2, Green: mode 3. Left column: goodness-of-fit analysis of regressions of mode frequencies vs syllabic rates obtained from the sound envelope. Right column: goodness-of-fit analysis of regressions of mode frequencies vs syllabic rates obtained from air sac pressure. Residuals are homoscedastic in all cases, p>0.05, Breusch-Pagan test, White test. **B, D, F, H, J, and L)** Normality tests for the residuals of the linear regression fit between mode frequencies and syllabic rates obtained from the envelope (panels B, D, and F) or air sac pressure (H, J, and L). *Top panel:* residual distribution histogram with normal distribution trace superimposed (black line). *Bottom panel:* quantile-quantile plot showing the empirical quantiles of the residuals (y-axis) as a function of the expected quantiles of a normal distribution with the same mean and variance as the empirical distribution (x-axis). Residuals are normally distributed in all cases (p>0.05, Shapiro-Wilks test, Anderson-Darling test).

**Supplementary Figure S12.**
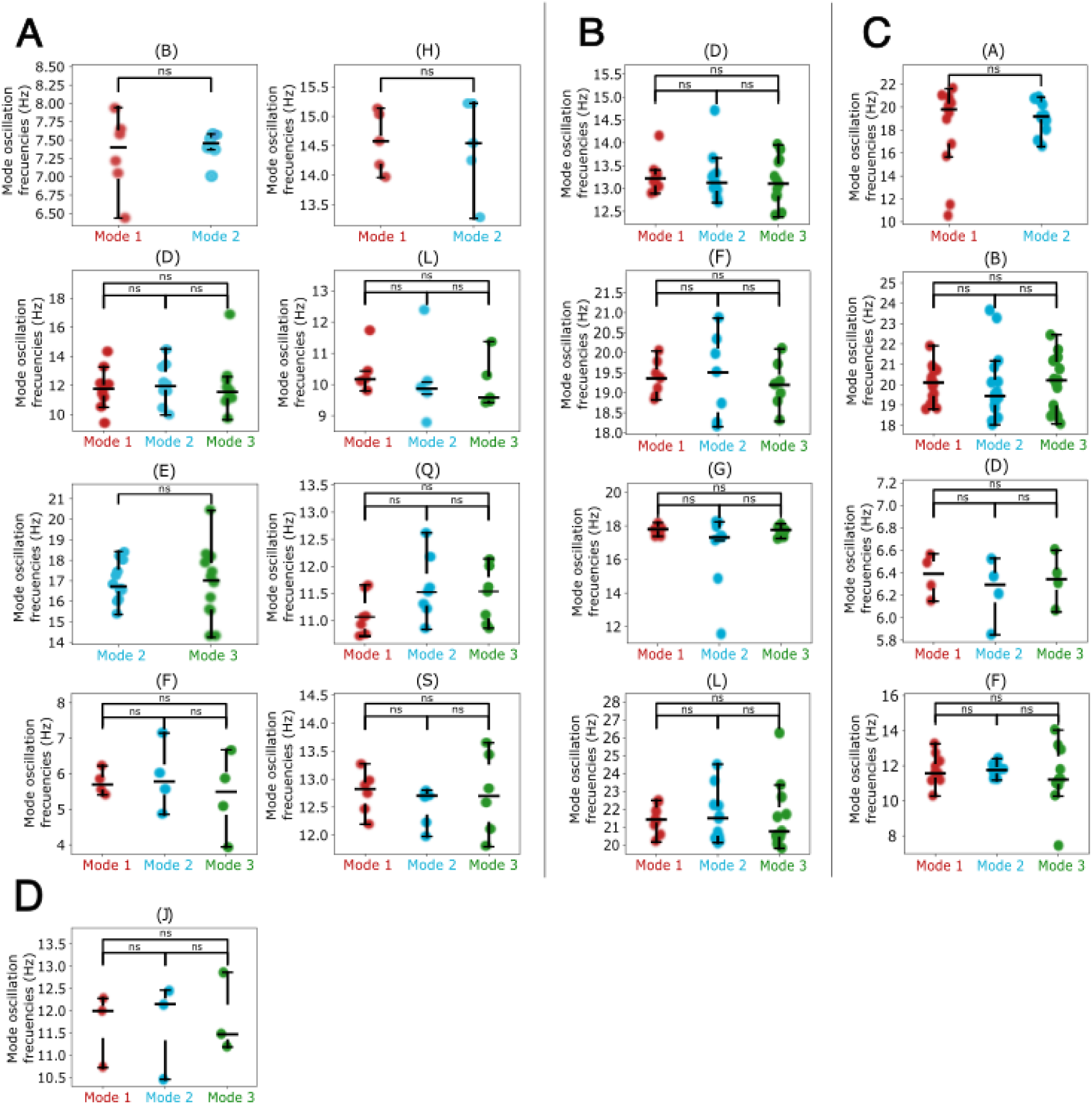
Distribution of frequencies calculated for mode oscillations in each phrase. All subplots show boxplots comparing the frequencies for mode 1 (red), mode 2 (light blue), and mode 3 (green) calculated for each phrase (indicated at the top of each subplot). In cases where data for any mode were discarded (for more details, see methods), only those used in the linear fits shown in figures 4 and supplementary S10 are displayed. Panel A shows the calculated frequencies for the phrases of caFLL001-VioAma, panel B the phrases of caCTH2447-RoNa, panel C the phrases of caCTH03-VioAma, and panel D those of caCTH140-VioVe. The boxplot of the phrases that have 2 to 3 syllables per phrase were omitted due to low data quantity (1-2 points per phrase). Frequency distributions show no significant differences between different modes (Kruskal-Wallis test, p>0.05); therefore, there is not enough evidence to assert that the data of the different modes come from different populations, so the frequency of the same phrase in different modes is similar.

## Notes

### Competing Interest Statement

The authors have declared no competing interest.

